# A biofidelic Goat Model of Traumatic Optic Neuropathy with Optic Canal Fracture via Transnasal Endoscopy

**DOI:** 10.64898/2026.02.18.705240

**Authors:** Zhonghao Yu, Hengzhuo Duan, Tonghe Yang, Yi Cao, Shudong Tian, Huan Wu, Jiale Zhang, Yue Wang, Ruixin Zhou, Shengjian Lu, Boyue Xu, Mengyun Li, Tian Xia, Si Zhang, Haodi Chen, Shurui Huang, Yikui Zhang, Jian Yang, Wencan Wu

## Abstract

Traumatic optic neuropathy (TON) is a major cause of irreversible vision loss following blunt cranial trauma, yet the absence of clinically relevant large animal models that faithfully recapitulate human TON has significantly impeded translational research. Current rodent models do not reproduce optic canal fracture, a key injury mechanism in many patients with TON. Here, we combined high-fidelity finite element analysis (FEA) with iterative engineering to establish a reproducible goat model of TON. We first built a high-resolution human-head finite element model to characterize force transmission to the optic nerve. Across clinically relevant periorbital impacts, stress preferentially concentrated in the intracanalicular segment, reaching a peak force density of approximately 500 N/m^2^ at 50.6 ms, about fivefold higher than in the intraorbital segment. Simulations further showed that direct optic canal impact reproduced comparable intracanalicular stress with a markedly lower input force: 195 N, compared with 3900 N for periorbital impact. Guided by these insights, we developed transnasal endoscopic impact systems capable of inducing optic canal fractures in goats. TON was confirmed within 24 hours by a characteristic relative afferent pupillary defect (RAPD), and at 1 month post-injury goats (n = 14) exhibited a 10%-20% reduction in ganglion cell complex (GCC) thickness and 40%-65% reductions in flash visual evoked potential (FVEP) and pattern electroretinogram (PERG) amplitude ratios (all P ≤ 0.001), with structural and functional preservation of the fellow eye. This study presents a robust, standardized, and clinically relevant large-animal platform for investigating TON pathophysiology.

## Introduction

Optic neuropathies represent a leading cause of irreversible blindness worldwide, posing a significant challenge to modern ophthalmology despite advances in our understanding of central nervous system regeneration (Laha et al., 2017; Lee et al., 2024). While recent studies have elucidated key intrinsic and extrinsic mechanisms regulating retinal ganglion cells (RGCs) survival and axon regeneration (Benowitz et al., 2017), the translation of these findings into clinical practice remains stalled (Tsai et al., 2024). A major barrier to bridging this translational gap is the absence of large-animal models that accurately recapitulate the anatomical, biomechanical, and pathological features of human optic neuropathy (Zhang et al., 2021). Moreover, most existing experimental paradigms are defined empirically rather than by quantitatively grounded cranio-orbital biomechanics, limiting standardization and cross-study comparability.

TON serves as a particularly valuable disease paradigm within the spectrum of optic neuropathies (Chen et al., 2022). Characterized by acute onset and a well-defined injury site (Yu-Wai-Man, 2015), TON shares pathological features with conditions like glaucoma yet offers a more precise temporal window for therapeutic intervention. Currently, TON research has relied heavily on rodent models. However, the rodent visual system differs fundamentally from that of humans in terms of optic canal morphology and skull architecture (Ibrahim et al., 2018; Tao et al., 2017). Furthermore, commonly used ultrasound-based or optic nerve crush models often lack the biomechanical fidelity of clinical optic canal fractures and induce widespread collateral damage to periorbital tissues (Tang et al., 2011; Tao et al., 2017). Together, these limitations highlight the need for a clinically grounded large-animal model that captures optic canal–related injury mechanics in a standardized and quantitative manner.

To address these limitations, we sought to establish a model in a species with closer anatomical similarity to humans. Our previous comparative anatomical studies identified the goat *(Capra hircus)* as an optimal candidate (Zhang et al., 2022), possessing an optic canal and sphenoid sinus structure highly similar to human anatomy and amenable to transnasal endoscopic access. In this study, we employed finite element analysis (FEA) to map stress distributions under periorbital trauma and identified localized optic canal impact as an efficient modeling strategy. We then engineered a standardized impact system and applied it transnasally under endoscopic guidance to induce optic canal fracture, thereby establishing a highly reproducible goat model of TON that closely recapitulates the clinical features of the human condition.

## Materials and methods

All materials used in the study are listed in Table 1.

**Table 1.** Materials and supplies.

| **Materials** | **Company** | **Catalogue no.** |
| --- | --- | --- |
| Saanen goat | Caimu Livestock Company,<br>China | - |
| Propofol | Fresenius Kabi, Sweden | - |
| Isoflurane | RWD Life Science, China | - |
| Xylazine | Jilin Huamu Animal Health<br>Products Co. LTD, China | - |
| periosteal elevator | JZ Classic, China | - |
| Nasal Mucosa Knives | JZ Classic, China | H7Z150 |
| Sinusscopes | HENKE SASS WOLF,<br>Germany | 7197200300 |
| Endoscopic<br>microdebrider | Medtronic, America | 1884004 |
| Endoscopic microdrill | Medtronic, America | 1882969 |
| Closed IV catheter<br>system | BD Shanghai Medical Device<br>Co., Ltd, China | 383019 |
| Hemocoagulase Atrox | Penglainuokang<br>Pharmaceutical, China | - |
| Heidelberg Spectralis<br>OCT system | Heidelberg Engineering<br>(Germany) |  |
| Electrophysiology<br>System | GOTEC Co., Ltd., China | GT-2008V-III |
| GraphPad Prism | GraphPad Prism | RRID:SCR_002798 |

### Animals

The male Saanen goats (4-7 months old) were purchased from Caimu Livestock Company (Hangzhou, China) and housed in the animal facility at Wenzhou Medical University. All experimental animals were housed in temperature-controlled rooms under a regular light–dark cycle, with unrestricted access to food and water. Animals were allowed a minimum acclimatization period of one week prior to the initiation of experiments. All experiments were approved by the Institutional Animal Care and Use Committee at Wenzhou Medical University (Wenzhou, China, ID number: wydw2022-0523). A total of 14 male Saanen goats were used (gas-driven impactor, n=8; elastic-energy impactor, n=6). In all animals, the left eye underwent optic canal impact and the contralateral right eye served as an internal control for longitudinal assessments.

### Finite element analysis

Finite element simulations were performed using COMSOL Multiphysics 6.0 to quantify the transient stress response of the optic nerve under head-impact loading. The workflow comprised CT-based geometry reconstruction, tetrahedral meshing, boundary condition settings, and time-dependent dynamic analysis under prescribed impact force.

### Geometry Reconstruction and Meshing

An anonymized head CT dataset (continuous slice thickness: 0.7 mm) was used to reconstruct the subject-specific geometry. The CT images were acquired using a SOMATOM go.Top CT scanner (Siemens Healthineers, Germany). The dataset was obtained with informed consent under an institutional review board–approved protocol (IRB no. 2024-065-K-57-03), in accordance with the Declaration of Helsinki. To improve computational efficiency, a simplified head model was constructed (Li et al., 2020), incorporating five key components: the optic nerve, skull, eyeball, extraocular muscles, and tendons. This model omitted complex structures such as the brain and blood vessels, including only five biological tissues: the unilateral optic nerve, skull, eyeball, extraocular muscles, and tendons. Intracranial soft tissues (e.g., brain parenchyma) were excluded to focus computational resources on the orbital region. Image segmentation was performed in 3D Slicer (v5.0.2). The skull was extracted via HU thresholding (200∼2000 HU) followed by post-processing steps including hole filling and island removal (Maire and Withers, 2014). The optic nerve, eyeball, extraocular muscles, and tendons were manually delineated based on anatomical landmarks. The segmented label maps were then converted into 3D surface meshes and processed using Laplacian smoothing (iterations = 30, *λ*= 0.5) to minimize staircase artifacts inherent to voxel-based reconstruction. Upon completion of geometric processing, the smoothed surface models were exported in Standard Tessellation Language (STL) format.

These STL files were then imported into COMSOL Multiphysics 6.0 for volumetric meshing and simulation. The domain was discretized using linear tetrahedral elements. The global element size range was set to 2.5∼14 mm, with adaptive local refinement applied near the optic nerve to a minimum size of 0.6 mm (growth rate = 1.5). The final mesh comprised 173,215 tetrahedral elements.

### Material properties

In this study, the modeling of all structures was conducted using isotropic and homogeneous material properties. The focus for elastic materials was placed on three fundamental mechanical parameters: Young’s modulus (*E*), Poisson’s ratio (𝜈), and density (*ρ*). Specifically, the skull was modeled using a linear elastic material model with a Young’s modulus of 14.5 GPa and a Poisson’s ratio of 0.35. Given the contrast in compressibility between tissues such as muscles and ligaments and the near incompressibility of the eyeball, the latter was modeled with a material having a Poisson’s ratio close to 0.5 (Li et al., 2020) to reflect its incompressible nature. Moreover, the biomechanical behavior of the optic nerve was represented using a linear viscoelastic material model widely adopted in numerical models, assuming that the time-varying shear modulus is independent of strain magnitude. Material parameters for head tissues were referenced from previous literature (Li et al., 2020), and a comprehensive table of material properties is provided in this document (Table 2).

**Table 2.** Material properties.

| tissue | density (kg/ m<sup>3</sup>) | Material parameters |
| --- | --- | --- |
| optic nerve | 1040 | $E_{\infty}=30\text{kPa}$, $\nu\sim 0.5$;<br>$g_1=0.45, \tau_1=0.5; g_2=0.365 \tau_2=50$ |
| skull | 1300 | $E=14.5\text{GPa}$, $\nu=0.35$ |
| eyeball | 1006 | $E=2.272\text{GPa}$, $\nu\sim 0.5$ |
| extraocular<br>muscles | 1200 | $E=11\text{MPa}$, $\nu=0.4$ |
| ligament | 1109 | $E=5\text{MPa}$, $\nu=0.4$ |

For the mathematical description of linear viscoelastic materials, the generalized Maxwell model was approximated using the Prony series. The linear viscoelastic model was expressed by the following equation:

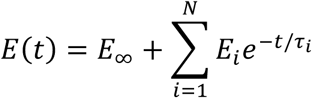

where *E*_∞_ represents the material’s stiffness at infinite time, *E*_*i*_ is the stiffness at the i time point, and *τ*_*i*_ is the corresponding relaxation time. The shear modulus *G*(*t*) was calculated as:

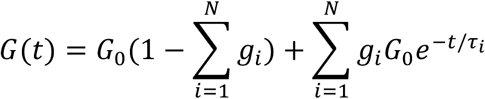

here, *g*_*i*_ = *G*_*i*_/*G*_0_, with *G*_0_ being the instantaneous shear modulus. Through this methodology, our study accurately simulated the complex behaviors of biological tissues under load and provided a reliable numerical method for advancing the understanding of biomechanical phenomena.

Detailed settings of material property parameters are shown in Table2.

### Boundary condition setting and analysis

Transient dynamic analyses were conducted using the solid mechanics module of COMSOL Multiphysics. This research was designed with reference to the simulation studies on the impact of cylindrical projectiles on the skull by Huempfner-Hierl (Huempfner-Hierl et al., 2015) and the demonstration by Chen (Chen and Ostoja-Starzewski, 2010) that the time course of the impact force during collisions approximates a Gaussian function. Impact loading was prescribed as a transient total force F(t) governed by a Gaussian function:

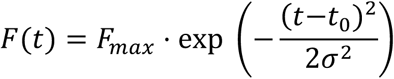

where *F*_*max*_= 3900 N, *t*_0_= 50 ms, and *σ*= 10 ms. To implement this load on the finite element mesh, the total force was applied as a uniformly distributed surface traction over an elliptical contact patch (Area = 1 cm^2^). Mathematically, the applied boundary traction *T*(*t*) was defined as the total force normalized by the contact area:

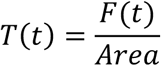

Fixed constraints were applied to nodes on the skull base. To investigate the sensitivity of the optic nerve to impact location, loads were applied to four quadrants of the orbital rim (superior, inferior, lateral, medial) and directly to the optic canal region. Directional sensitivity was assessed by perturbing the loading vector by ±10° relative to the surface normal. Additionally, a parametric study was conducted by varying the skull’s elastic modulus, Poisson’s ratio, and density by ±10%.

All simulations were performed using the transient solver in the solid mechanics module. Time integration was executed using the Generalized-α scheme to ensure optimal control of high-frequency numerical dissipation while maintaining accuracy for inertial dynamics. A constant time step of 1.1 ms was selected for the analysis. This temporal resolution was determined based on the impact duration (*t*_0_= 50 ms) to ensure sufficient sampling density. This resolution allowed for the accurate capture of both the Gaussian force profile and the time-dependent viscoelastic relaxation of the optic nerve, while maintaining computational efficiency. The total simulation time was set to 200 ms to observe the complete stress evolution from impact initiation to relaxation. Finally, to assess the clinical relevance of the computational results, a face validity evaluation was conducted. The simulated stress distribution patterns were qualitatively compared with typical injury locations observed in clinical cases by experienced neuro-ophthalmologists. This qualitative comparison served to confirm that the model predicts stress concentrations in anatomically plausible regions consistent with real-world trauma.

### Transnasal endoscopic procedure to expose the optic canal in goats

The procedure was described in detail in our previous study (Zhang et al., 2022), and is further refined in the present work. General anesthesia was induced by an intravenous bolus of propofol (10 mg/kg), followed by endotracheal intubation using a 6.0-mm cuffed tube (Henan Tuoren Medical Device Co., Ltd., China). Anesthesia was maintained with 2.5–3.0% isoflurane delivered in a 1:1 mixture of oxygen and air at a flow rate of 2 L/min via a mechanical ventilator. Vital signs, including heart rate, respiratory rate, and oxygen saturation, were continuously monitored throughout the procedure using an electrocardiographic monitoring system.

To reduce intraoperative bleeding, hemocoagulase atrox was administered intravenously (1–2 U per goat) after anesthesia induction. Prophylactic antibiotics and anti-inflammatory agents were administered intravenously, including gentamicin (4 mg/kg), ceftiofur sodium (0.1 g/10 kg), and dexamethasone sodium phosphate (1 mg/kg). The corneas were protected with carbomer gel to prevent exposure-related drying.

The surgical field extended from the forehead to the nasal dorsum, with the lateral boundaries defined by the medial canthi. Hair was shaved, and the skin was disinfected three times using 5% povidone–iodine solution (Zhejiang Apeloa Inc., China). Sterile surgical drapes were applied, leaving only the nasal bridge exposed.

A midline facial incision (approximately 4–5 cm) was first made at the level of the medial canthi, followed by a vertical incision along the nasal midline (approximately 4–5 cm) and an additional inferior horizontal incision of similar length. The periosteum and underlying bone were carefully dissected using toothed forceps and a periosteal elevator. The exposed nasal bones was removed to gain access to the nasal cavity.

Upon entering the nasal cavity, an endoscopic microdebrider equipped with a straight-cut microdebrider (Medtronic, Cat. #1884004) was used to partially resect the middle turbinate and posterior olfactory mucosa on the right side, allowing exposure of the anterior bony wall of the sphenoid body. The nasal septum was then incised using a falciform knife, and partial resection of the contralateral olfactory mucosa was performed to further enlarge the operative corridor and improve visualization.

Because goats lack a naturally pneumatized sphenoid sinus anterior to the optic canal, an artificial sphenoid sinus was created. A diamond microdrill (Medtronic, Cat. #1882969) was used to thin and remove the cortical bone overlying the central anterior wall of the sphenoid body. Drilling was performed under continuous endoscopic visualization until a characteristic loss of resistance (“give-way” sensation) was encountered, indicating entry into the sphenoid cavity. Residual bone debris was removed using a blunt dissector, and hemostasis was achieved with calcium alginate packing when necessary.

The bony window of the sphenoid cavity was then gradually enlarged to obtain a clear surgical field. After removal of the packing material, the bony landmarks of the prechiasmatic region and the anterior wall of the optic canal became clearly visible. Using a combination of coarse and fine diamond burrs, the anterior bony wall of the optic canal was carefully thinned and removed until the optic canal was fully exposed, providing direct access to the intracanalicular segment of the optic nerve.

### Modeling optic canal fracture in goats

The optic canal was identified at the junction of the anterior wall of the optic chiasm and the medial orbital wall under transnasal endoscopic visualization. The impactor tip was positioned in firm contact with the anterior bony wall of the optic canal, and the device was calibrated to deliver a predefined impact force. Upon activation, the metal impact rod delivered a focal mechanical strike to the optic canal wall, resulting in a localized fracture with impinged bone that was immediately visualized endoscopically.

This impact consistently generated a circular bony fragment approximately 2 mm in diameter, which became displaced into the optic canal and produced sustained compression of the intracanalicular optic nerve. Following completion of the impact procedure, the nasal cavity was irrigated and disinfected under sterile conditions. The artificially created sphenoid sinus was packed with an absorbable gelatin sponge to achieve hemostasis, and the surgical incision was closed using 3-0 sutures. Animals were then carefully recovered from general anesthesia.

Pupillary light reflex (PLR) testing was performed prior to surgery and again 24 h post-procedure to verify successful induction of TON. Loss of PLR was consistently observed in the injured eye, while normal PLR responses were preserved in the contralateral eye.

### Modified impact device and stabilizing base

To enhance the stability, reproducibility, and translational applicability of the optic canal impact model, the original high-pressure gas–driven impactor was redesigned. The conventional gas-driven system is inherently susceptible to pressure fluctuations and potential gas cylinder leakage, both of which can compromise impact consistency and experimental reproducibility. In the modified device, a motor-controlled mechanism was introduced to quantitatively stretch a compression spring, which subsequently releases stored elastic energy to propel the impact head. The spring used in this system (1.5 × 12 × 45 mm) was calibrated such that a stretching length of 20 mm produced an impact effect comparable to that of the original gas-driven device. By changing the mode of energy delivery from gas-driven to elastic energy–driven propulsion, this elastic energy–driven impactor provided improved stability and reliability for large-animal modeling.

During optic canal fracture induction in goats, animals were positioned prone, allowing the operator to face the animal’s head directly and enabling handheld use of the impact device. In contrast, nonhuman primates are typically operated in the supine position, rendering handheld impact delivery impractical and potentially unstable. To further enhance impact precision and extend the applicability of this model across species, we developed a dedicated stabilizing base for the impactor. This base incorporated five independently driven motors, which enabled multi-axis adjustment to accommodate different surgical orientations, minimize recoil, and eliminate positional deviations associated with handheld operation. Collectively, these modifications substantially improved the precision and reproducibility of optic canal fracture induction and provide a scalable platform for translational studies of TON in large-animal and nonhuman primate models.

### Non-invasive visual function assessment in goats

Visual function in goats was evaluated using pupillary light reflex (PLR), optical coherence tomography (OCT), flash visual evoked potentials (FVEP), and pattern electroretinography (PERG). All procedures were performed according to standardized protocols refined from our previous work (Zhang et al., 2022), with detailed descriptions provided below.

For all examinations, endotracheal intubation was performed under laryngoscopic guidance to ensure airway patency and minimize the risk of aspiration.

For PLR, PERG, and OCT examinations, goats were sedated by intramuscular injection of xylazine (3 mg/kg). For FVEP recordings, general anesthesia was induced with intravenous propofol, followed by maintenance with inhaled isoflurane (2.5–3.0%) delivered at a flow rate of 2 L/min via endotracheal intubation.

#### PLR

Prior to PLR testing, animals underwent 5 min of dark adaptation to stabilize baseline pupil diameter. Eyelid specula were placed bilaterally to fully expose the globes, and artificial tears were applied to maintain corneal hydration throughout the procedure.

PLR measurements were obtained using a custom-built PLR recording system connected to a synchronized video acquisition unit and programmable light source. The camera was positioned to simultaneously capture both eyes and adjusted to achieve stable, high-resolution imaging. Recordings were performed under both visible light and infrared illumination to enable continuous, real-time monitoring of pupil dynamics.

During testing, a white-light stimulus (230 lx, 2 s duration) was delivered monocularly to elicit pupillary constriction, followed by a 12 s interval to allow full redilation. This stimulation sequence was repeated three times for each eye. The non-stimulated eye was occluded during testing to prevent consensual responses from confounding measurements.

PLR responses were graded semi-quantitatively based on pupillary reactivity: Grade 0 indicated complete absence of response; Grade 1 indicated a reduced response characterized by delayed constriction and decreased amplitude; Grade 2 indicated a brisk and robust constriction response.

#### OCT

For OCT imaging, topical anesthesia was achieved by instillation of proparacaine hydrochloride eye drops, followed by pharmacological mydriasis using tropicamide. Eyelid specula were applied to maintain full exposure of the pupil.

Retinal imaging was performed using a Heidelberg Spectralis OCT system (Heidelberg Engineering, Germany). The optic nerve head region was centered along the optical axis of the instrument, and peripapillary scans were acquired using a circular scanning protocol. For each eye, 100 consecutive high-resolution scans were obtained from the same retinal location and averaged using the system’s built-in software to generate final images with enhanced signal-to-noise ratio. GCC thickness was quantified for subsequent analysis.

#### PERG

Prior to PERG recording, topical proparacaine hydrochloride and tropicamide eye drops were administered, and eyelid specula were placed bilaterally. The frontal hair was shaved, and the skin was disinfected three times using alternating povidone–iodine and 75% ethanol.

A small midline incision was made in the frontal skin, and a sterilized screw electrode was inserted into the frontal bone to serve as the ground electrode. The screw was secured using an alligator clip to ensure signal stability. Needle electrodes were placed approximately 1 cm lateral to each outer canthus as reference electrodes. ERG-Jet corneal electrodes, pre-coated with carbomer gel, were gently positioned on the center of each cornea as recording electrodes.

Electrode impedance was verified to be below 10 kΩ prior to data acquisition. PERG signals were recorded using a commercial electrophysiology system (GT-2008V-III, GOTEC Co., Ltd., China). Visual stimuli were presented on two 22-inch monitors (1920 × 1080 resolution) positioned approximately 50 cm in front of each eye. The stimulus consisted of contrast-reversing black-and-white checkerboard patterns with a temporal frequency of 2.4 Hz, contrast of 96%, and mean luminance of 200 cd/m². Recordings were performed at spatial frequencies of 0.1, 0.3, 3.0, and 12.6 cycles per degree (cpd).

Signals were amplified 16,000-fold, band-pass filtered between 1 and 100 Hz, and averaged over 64 consecutive sweeps. In the averaged waveform, the first positive peak was designated as P1 (typically ∼25 ms), and the first negative peak as N1 (typically ∼55 ms). PERG amplitude was measured from N1 to P1. Functional outcomes were expressed as the amplitude ratio between the injured eye and the contralateral eye, with greater reductions indicating more severe RGC dysfunction.

#### FVEP

FVEP recordings were performed under general anesthesia using procedures identical to those described for PERG with respect to intubation, ocular preparation, hair removal, and skin disinfection. Sterilized screw electrodes were implanted in the central frontal bone and central occipital bone to serve as reference and recording electrodes, respectively. A needle electrode placed subcutaneously near the reference electrode served as the ground. Electrode impedances were confirmed to be below 10 kΩ before recording.

Animals underwent 5 min of dark adaptation prior to testing. The tested eye was held open using an eyelid speculum, and flash stimuli of three intensities (0.025, 0.25, and 3.0 cd·s/m²) were delivered under scotopic conditions. Each stimulus intensity was recorded twice, and the averaged waveform was used for analysis. A 2 min dark adaptation interval was provided between stimulus conditions, during which artificial tears were applied to maintain corneal hydration.

Both eyes were tested separately, and the non-tested eye was occluded throughout the procedure to prevent binocular interference.

### Definition of TON and severity assessment

The diagnosis of TON was based on clinical functional criteria indicative of afferent visual pathway dysfunction in the injured eye. Specifically, TON was defined by the presence of a relative afferent pupillary defect (RAPD), operationally identified as an inter-eye difference in pupillary light reflex (PLR) grade of ≥2 within 24 h after impact (injured eye PLR grade = 0, while the contralateral eye maintained a grade of 2). Goats that did not meet this predefined criterion were excluded from subsequent analyses.

The extent and severity of TON were subsequently evaluated using continuous structural and functional outcome measures, including: (i) the percentage reduction in retinal GCC thickness of the injured eye at 1 month post-injury (1 mpi) relative to both baseline values and the contralateral eye; (ii) the percentage decrease in flash visual evoked potential (FVEP) P1–N1 amplitude ratio (injured eye/contralateral eye) relative to baseline; and (iii) the percentage decrease in pattern electroretinography (PERG) P1–N1 amplitude ratio (injured eye/contralateral eye) relative to baseline. These quantitative indices were used for statistical analyses to compare differences across experimental groups and time points, with greater percentage reductions indicating more severe TON.

### Statistical analysis

Statistical analyses were performed using GraphPad Prism 9.0 (GraphPad Software, San Diego, CA, USA) and R software (R Foundation for Statistical Computing, Vienna, Austria). All data are presented as mean ± SEM. Normality and homogeneity of variance were assessed using the Shapiro–Wilk and Levene tests, respectively. Depending on whether these assumptions were met, data were analyzed using ordinary one-way ANOVA, two-way ANOVA, or the Scheirer– Ray–Hare non-parametric two-factor test, as specified in the figure legends. Post hoc comparisons were performed using Tukey’s test or appropriate non-parametric multiple comparison procedures. A p value < 0.05 was considered statistically significant.

## Results

### Finite element analysis identifies the intracanalicular segment as the primary site of stress concentration

TON is a major cause of sudden and severe visual loss following blunt head trauma (Crompton, 1970; Gise et al., 2018), and the optic canal has long been implicated as the principal site of injury due to force transmission along the orbital wall (Fig. 1A)(Li et al., 2020; Wu et al., 2015). Consistent with this concept, optic canal fractures were observed in approximately 70% of TON patients in our previous multicenter clinical study (Wu et al., 2015).

**Figure 1.**
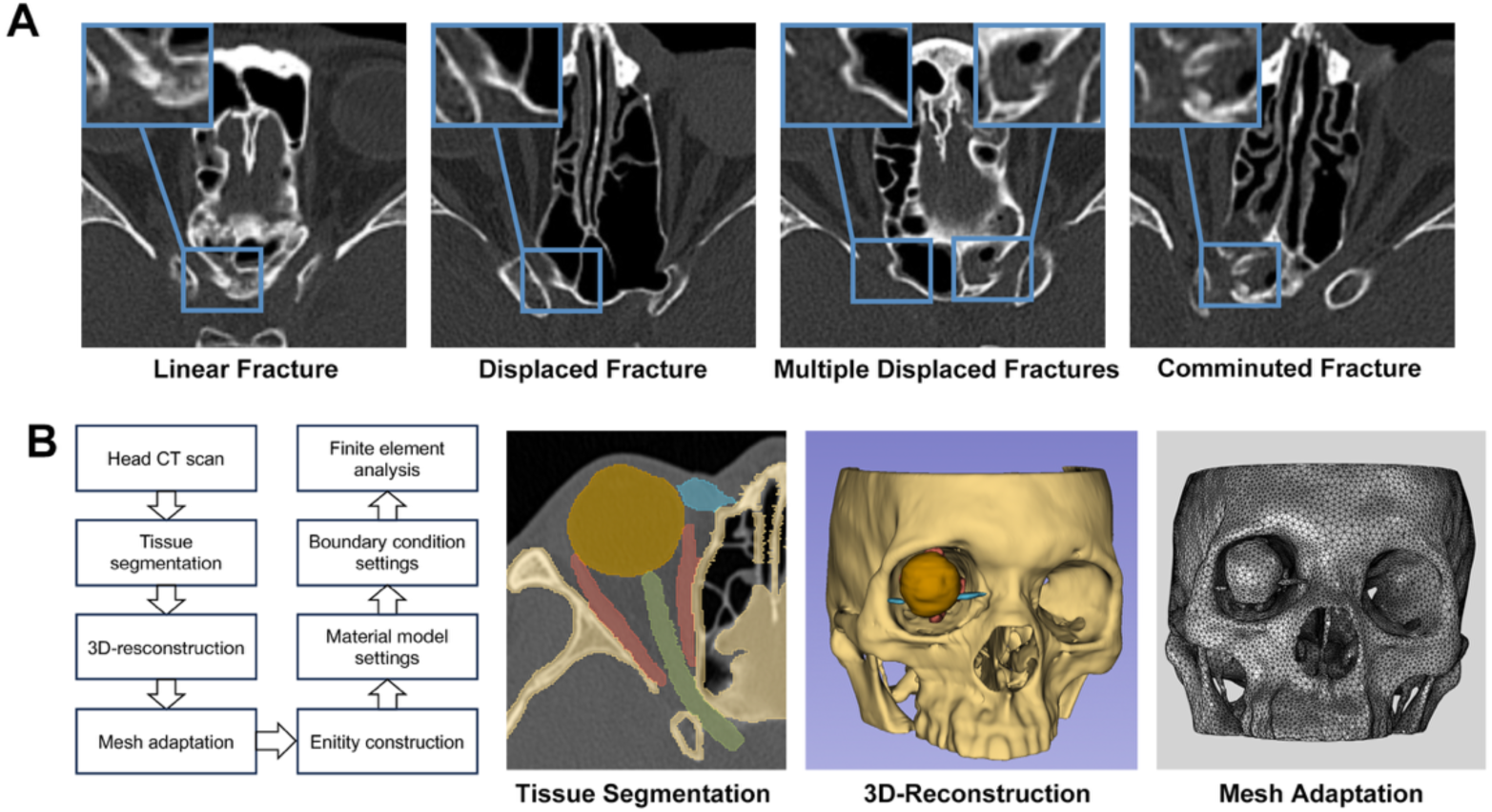
Finite element model of the human head used to simulate force transmission to the optic canal. **(A)** Representative horizontal head CT images depicting various optic canal fractures in clinical TON cases. **(B)** Flowchart and representative images demonstrating finite element analysis based on head CT scan step by step, including tissue segmentation, 3D-reconstruction, and mesh adaptation.

To elucidate the biomechanics of force transmission along the optic nerve, we constructed a high-fidelity finite element model based on high-resolution human skull CT scans (Fig. 1B). We simulated a clinically relevant temporal orbital wall impact using a force of 3900 N with a Gaussian distribution (Fig. 2A). Force distribution demonstrated that impact-induced forces propagated along the orbital wall and converged toward the optic canal region (Fig. 2B).

**Figure 2.**
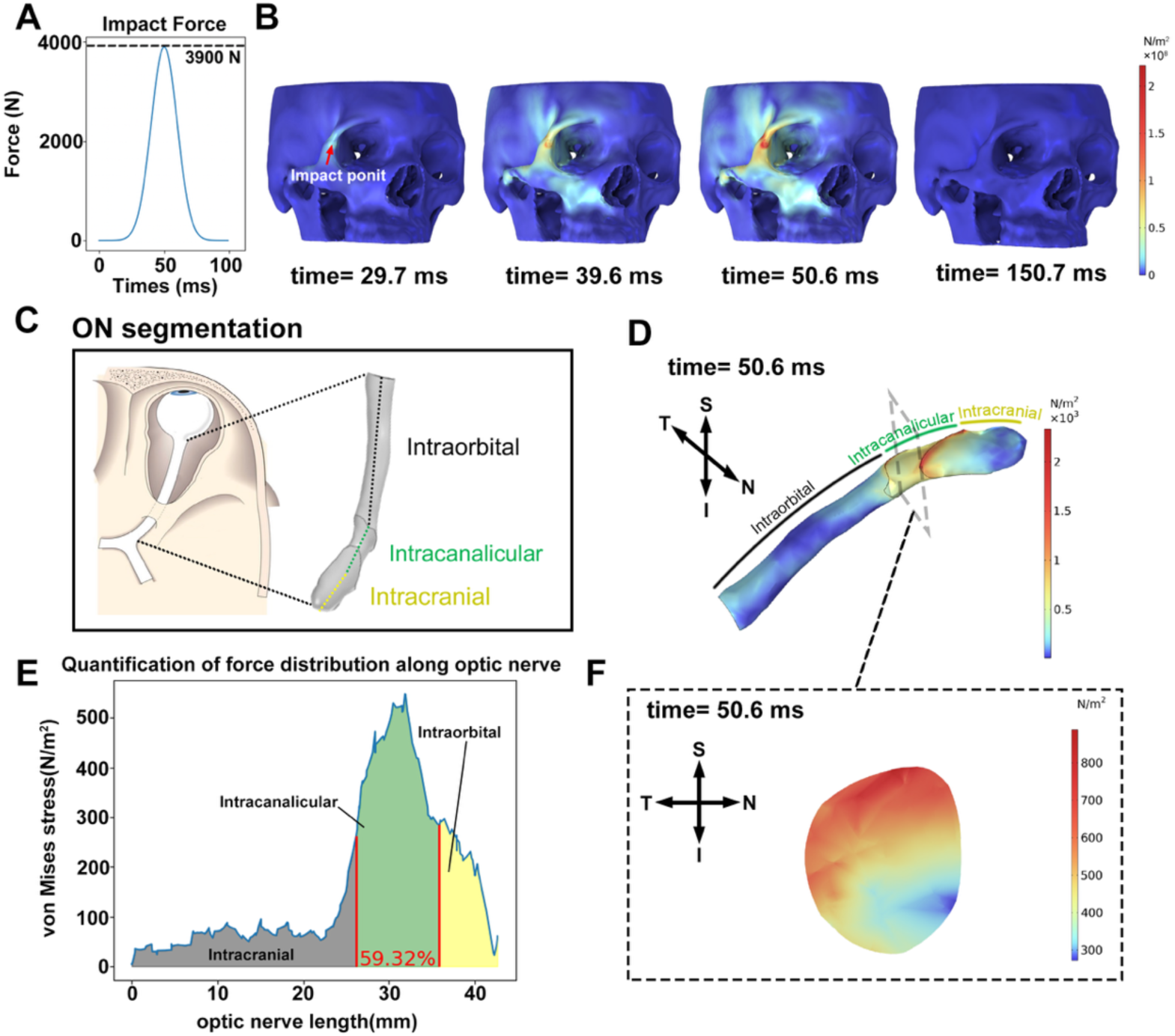
The intracanalicular optic nerve suffered the highest mechanical stress following a periorbital blunt impact. **(A)** Impact force reached its peak at 50 ms, exhibiting a Gaussian distribution with a magnitude of 3900 N. **(B)** Color-coded displays showing force propagation from the periorbital region towards the optic canal along the orbital wall. **(C)** Anatomic segmentation of the optic nerve (ON). **(D-E)** Color-coded illustration and quantification of force distribution along the ON at 50.6 ms. **(F)** Cross-sectional stress map showing the force distribution at the center of the intracanalicular segment of the optic nerve at 50.6 ms.

For quantitative analysis, the optic nerve was anatomically subdivided into intraorbital, intracanalicular, and intracranial segments (Fig. 2C). Following temporal periorbital impact, mechanical stress along the optic nerve central axis was preferentially concentrated within the intracanalicular segment (Fig. 2 D–F). Specifically, peak force density in the intracanalicular segment reached approximately 500 N/m² at 50.6 ms, which was nearly fivefold higher than that observed in the intraorbital segment (∼100 N/m² at the same time point). Notably, comparable stress concentration patterns were consistently observed in simulations of superior, inferior, and nasal periorbital impacts (Fig. S1), collectively identifying the intracanalicular segment as the primary biomechanical weak point during blunt orbital trauma.

### Direct optic canal impact caused little collateral injury and showed robustness against bias

Given the finite element analyses identifying the intracanalicular segment as the primary biomechanical vulnerability, two injury paradigms were considered for modeling TON: (i) indirect periorbital impact, in which external blunt force is transmitted to the optic canal through the orbital bones; and (ii) direct impact targeted to the optic canal itself. To determine which strategy provides superior safety and mechanical robustness, finite element analysis was used to compare these two approaches.

Simulations of indirect periorbital impact revealed widespread dispersion of mechanical forces across the maxillary bone, nasal bone, and skull base (Fig. 3A,B).

**Figure 3.**
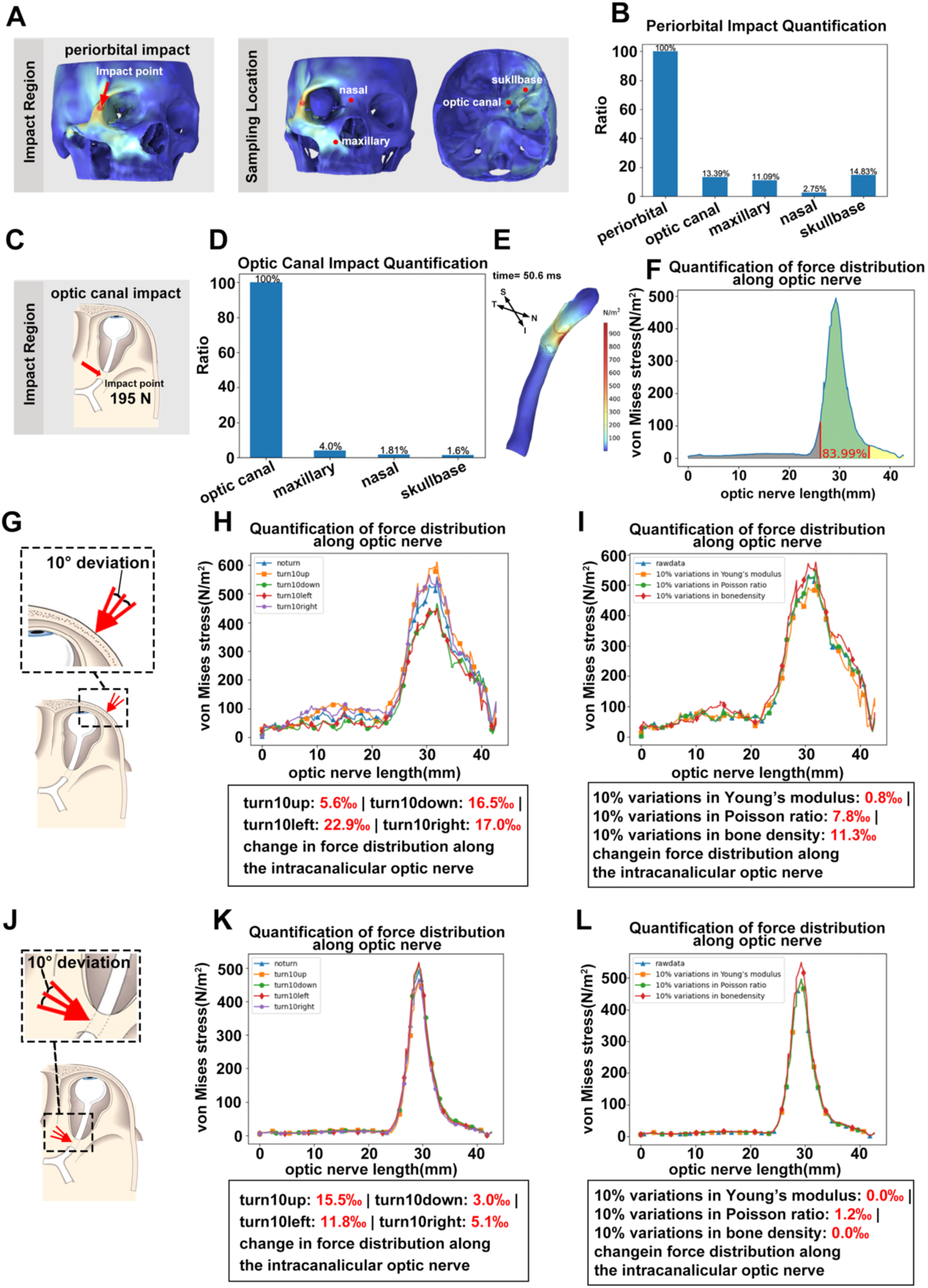
Computer simulation demonstrates that the optic nerve within the optic canal experiences the greatest mechanical stress after a clinically-relevant periorbital blunt impact. **(A)** Representative simulations illustrating the site of periorbital impact and the locations for force distribution analysis. **(B)** Quantification of force distribution at different analysis sites following periorbital impact at 50.6 ms. **(C)** Schematic showing direct impact to the optic canal. **(D)** Quantification of force distribution across different analysis sites following direct optic canal impact at 50.6 ms. **(E-F)** Color-coded illustration and quantification of force distribution along the optic nerve following direct optic canal impact at 50.6 ms. **(G, J)** Schematics illustrating deviation in impact angle by 10 degrees, both upward and downward, for periorbital (G) and optic canal (J) impacts. **(H-I, K-L)** Quantification of force distribution fluctuation along the intracanalicular optic nerve by 10-degree deviations in impact direction and a 10% variation in bone mechanical properties (density, Young’s modulus, Poisson’s ratio) for periorbital impact (H, I) and optic canal impact (K, L).

In contrast, direct optic canal impact caused minimal force propagation to adjacent skull structures (Fig. 3C,D), while producing a force distribution pattern along the optic nerve comparable to that induced by periorbital impact (Fig. 3E,F). Notably, direct optic canal impact required a substantially lower input force of 195 N, corresponding to approximately 5% of that used for temporal periorbital impact (3900 N), to achieve a similar peak force density within the intracanalicular segment (Fig. 3F), thereby reducing the likelihood of off-target skeletal injury.

Importantly, direct optic canal impact also demonstrated greater robustness against bias than indirect periorbital impact. Specifically, compared with indirect periorbital impact, a 10° deviation from the optimal vertical impact angle (Fig. 3G,H,J,K), as well as a 10% variation in bone mechanical properties, including density, Young’s modulus, and Poisson’s ratio (Fig. 3I,L), resulted in only minor fluctuations in force distribution along the intracanalicular optic nerve following direct optic canal impact.

Collectively, these results demonstrated that direct optic canal impact induces optic canal– centered injury with minimal collateral damage and high robustness against bias.

### A reproducible goat TON model with optic canal fracture is established by direct optic canal impact under transnasal endoscopy

Based on the finite element analysis results, we identified direct optic canal impact as a more robust injury paradigm and first engineered a gas-driven impactor to experimentally recapitulate TON. This device employed high-pressure air to propel an impact rod forward by 2 mm, thereby inducing an optic canal fracture with impinged bone (Fig. 4A–C). The air pressure within the storage chamber was manually adjustable from 1 to 10 megapascals (MPa), with higher pressures generating greater impact forces (Fig. 4B). An operating pressure of 3 MPa was selected, as it produced more consistent impact forces compared with lower (2 MPa) or higher (4 MPa) pressure settings (Fig. 4B).

**Figure 4.**
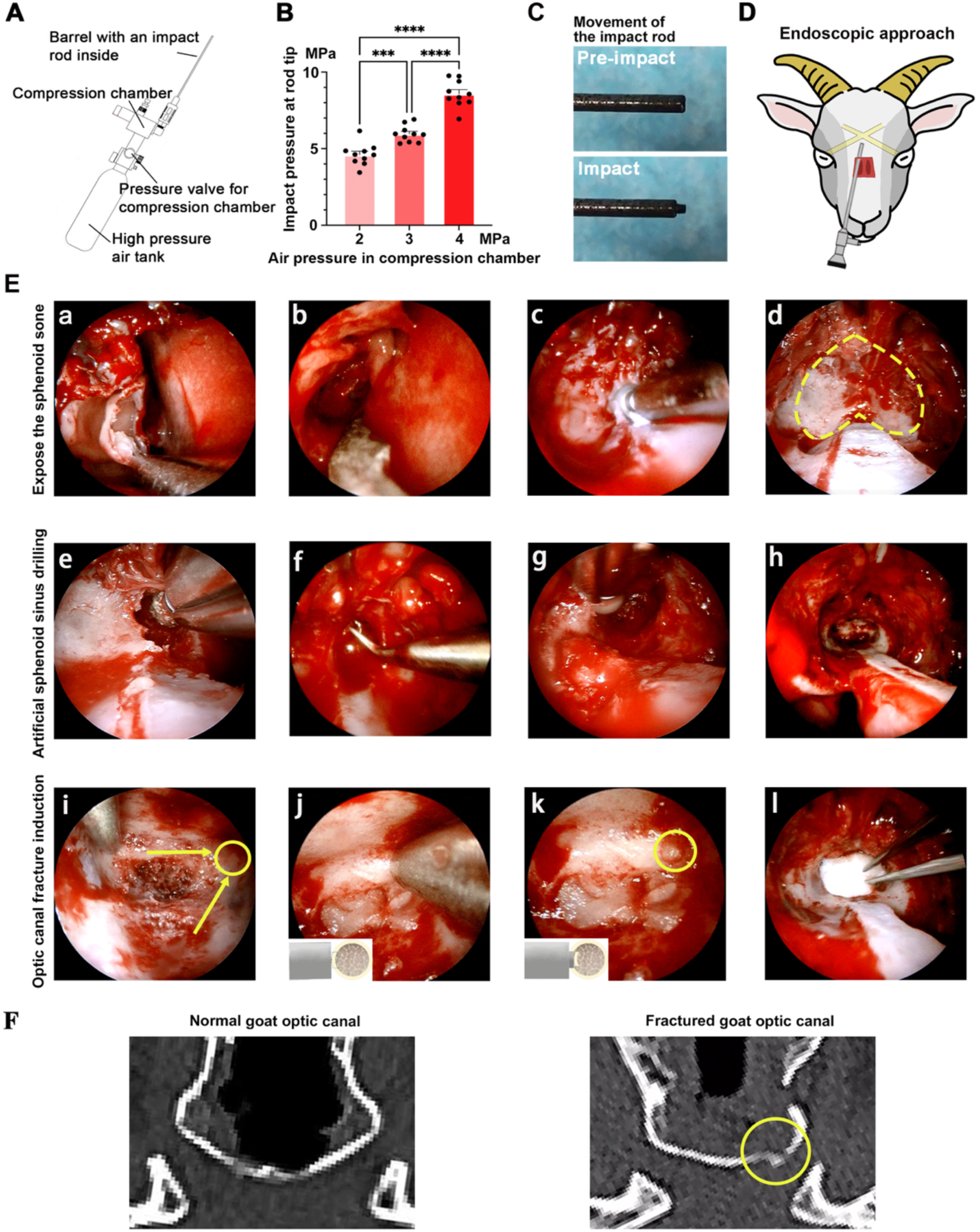
Establishment of a reproducible TON goat model induced by optic canal fracture under transnasal endoscopy. **(A)** Schematic diagram of the gas-driven impactor, consisting of a high-pressure air tank, a compression chamber with a pressure valve, and a barrel containing the impact rod. **(B)** Calibration of the device showing impact force at the rod tip under different air pressures in the compression chamber (2, 3, and 4 MPa). Each dot represents one impact (n = 10 per pressure level); data are shown as mean ± SEM. **(C)** Schematic diagram of the impact rod. **(D)** Schematic diagram of the endoscopic approach. **(E)** Intraoperative endoscopic views of key surgical steps. (a–d) Exposure of the anterior wall of the sphenoid bone after removal of the nasal septum, turbinates, and olfactory mucosa. The dotted yellow line in (d) outlines the bony window. (e–h) Drilling of the sphenoid body to create an artificial sphenoid sinus, providing direct access to the pre-chiasmatic optic canal. (i– l) Optic canal fracture induction: the impactor tip is seated against the anterior wall of the optic canal and a single impact generates a circumscribed bony fragment (yellow circle) that is displaced into the canal lumen. **(F)** Representative CT images showing a normal goat optic canal (left), a post-impact goat optic canal with a depressed fracture fragment (middle, yellow circle), and a human traumatic optic canal fracture for comparison (right, blue arrow).

Goats were chosen as the experimental species due to their human-like sinonasal anatomy, which facilitates minimally invasive transnasal endoscopic manipulation of the optic canal, together with their relatively favorable cost and availability compared with other large-animal models (Zhang et al., 2022). Under endoscopic visualization, the middle turbinate, nasal septum, and olfactory nerve bundles were sequentially resected using a straight-cut microdebrider or scissor (Fig. 4E-a,b,c), thereby exposing the anterior wall of the sphenoid bone, which exhibited a characteristic inverted heart-shaped appearance and served as a key anatomical landmark (Fig. 4E-d). The anterior cortical bone of the sphenoid was then carefully drilled to open the sphenoid cavity and create an artificial sphenoid sinus (Fig. 4E-e,f,g,h). Further bone removal within the artificial sphenoid sinus was performed using a diamond microdrill, which progressively widened the surgical corridor and allowed clear visualization of the optic canal within the operative field (Fig. 4E-i). The impact device was subsequently advanced along the predefined trajectory and precisely positioned against the anterior wall of the optic canal under endoscopic guidance (Fig. 4E-j). Upon triggering the impactor, a displaced, impacted optic canal fracture with inwardly displaced bone fragments was immediately observed under endoscopic visualization, resulting in direct compression of the intracanalicular optic nerve (Fig. 4E-k). Follow-up computed tomography (CT) imaging further confirmed that the fracture morphology closely resembled that observed in patients with TON (Fig. 1A, 4F).

Structural damage to retinal ganglion cells was quantified by OCT. In the injured eyes, peripapillary ganglion cell complex (GCC) thickness decreased from 244.5 ± 20.62 μm at baseline to 218.1 ± 17.99 μm at 1 month post-injury (1 mpi), corresponding to a ∼10.8% reduction (P = 0.0018, two-way ANOVA, time factor; Fig. 5A,B), whereas GCC thickness in the contralateral eyes remained unchanged (Fig. S2A), indicating preservation of retinal structure in the fellow eye.

**Figure 5.**
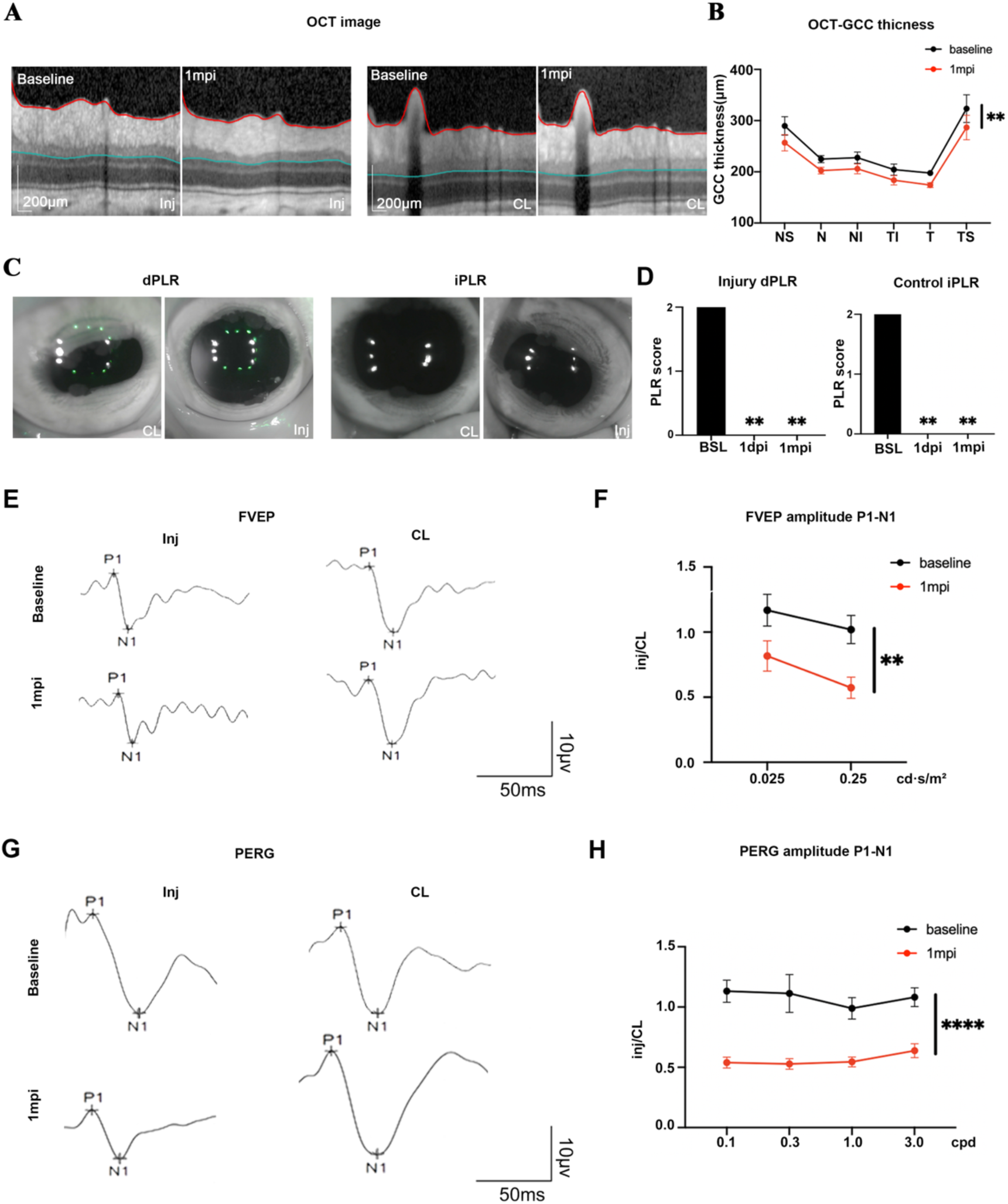
Structural and functional damage in a gas-driven goat model of TON. **(A)** Representative peripapillary OCT scans of the injured eye at baseline and 1 month post-injury (1 mpi), with the GCC segmentation overlaid. **(B)** Quantification of GCC thickness in different macular sectors at baseline and 1 mpi (n = 8 goats; mean ± SEM). In the injured eyes, GCC thickness decreased from 244.5 ± 20.62 μm at baseline to 218.1 ± 17.99 μm at 1 mpi (∼10.8% reduction; P = 0.0018, two-way ANOVA, time factor). **(C)** Representative infrared images of the direct (dPLR) and indirect (iPLR) pupillary light reflexes in the injured and contralateral eyes at 1 month post-injury (1 mpi). **(D)** Quantification of PLR scores for the contralateral eye dPLR (top graph) and injured eye iPLR (bottom graph) at baseline (BSL), 1 day post-injury (1 dpi), and 1 month post-injury (1 mpi). Data are shown as mean ± SD; ns, not significant. **(E)** Representative flash visual evoked potentials (FVEP) recorded from injured and contralateral eyes at baseline and 1 mpi. **(F)** Ratios of FVEP P1–N1 amplitudes between injured and contralateral eyes (Inj/CL) at baseline and 1 mpi (n = 8; mean ± SEM). The FVEP amplitude ratio decreased from 1.094 ± 0.07412 at baseline to 0.6950 ± 0.1217 at 1 mpi (∼36.5% reduction; P = 0.0010). **(G)** Representative pattern electroretinogram (PERG) traces from injured and contralateral eyes at baseline and 1 mpi. **(H)** Ratios of PERG P1–N1 amplitudes between injured and contralateral eyes (Inj/CL) at different spatial frequencies at baseline and 1 mpi (n = 8; mean ± SEM). The PERG amplitude ratio declined from 1.079 ± 0.03150 at baseline to 0.5626 ± 0.02553 at 1 mpi (∼47.9% reduction; P < 0.0001 at 0.3 cpd). OCT-based GCC thickness in (B) was analyzed using two-way ANOVA with retinal sector and time (baseline vs 1 mpi) as factors. PLR data in (D) were analyzed using one-way ANOVA. FVEP amplitude ratios in (F) were analyzed using two-way ANOVA with stimulus frequency and time as factors. PERG amplitude ratios in (H) were analyzed using the Scheirer–Ray–Hare non-parametric two-factor test with spatial frequency and time as factors. *p < 0.05, **p < 0.01, ***p < 0.001, ****p < 0.0001.

Functionally, the direct pupillary light reflex (dPLR) in the injured eye and the indirect PLR (iPLR) in the contralateral eye were both markedly reduced over time (Fig. 5C,D). In contrast, the dPLR in the contralateral eye and the iPLR in the injured eye remained intact (Fig. S2B). This dissociated PLR pattern met the predefined criteria for a relative afferent pupillary defect (RAPD) (Soleimani et al., 2011). Electrophysiological assessments showed consistent functional deficits in the injured eyes. The FVEP P1–N1 amplitude ratio (injured/contralateral) decreased from 1.094 ± 0.07412 at baseline to 0.6950 ± 0.1217 at 1 mpi (≈36.5% reduction; P = 0.0010; Fig. 5E,F). Similarly, the PERG P1–N1 amplitude ratio (injured/contralateral) declined from 1.079 ± 0.03150 to 0.5626 ± 0.02553 (≈47.9% reduction; P < 0.0001; Fig. 5G,H). In the contralateral uninjured eyes, FVEP and PERG amplitudes did not show significant changes compared with baseline (Fig. S2C,D).

Together, these results showed that transnasal endoscopic direct optic canal impact produced optic canal fracture and unilateral structural and functional deficits in goats. We then implemented a second-generation elastic-energy driven device for subsequent experiments.

### Elastic-energy–driven impact system for optic canal fracture modeling

To this end, we developed a second-generation elastic-energy driven, spring-based impact device for inducing optic canal fracture with impinged bone (Fig. 6A). The system employs a compression spring with dimensions of 1.5 × 12 × 45 mm, in which the stretching length is positively correlated with the delivered impact force (Fig. 6B). Based on mechanical calibrations, a deformation length of 20 mm was selected to generate an impact force equivalent to that produced by the gas-driven device at 3 MPa, thereby maintaining comparable injury severity while eliminating pneumatic instability.

**Figure 6.**
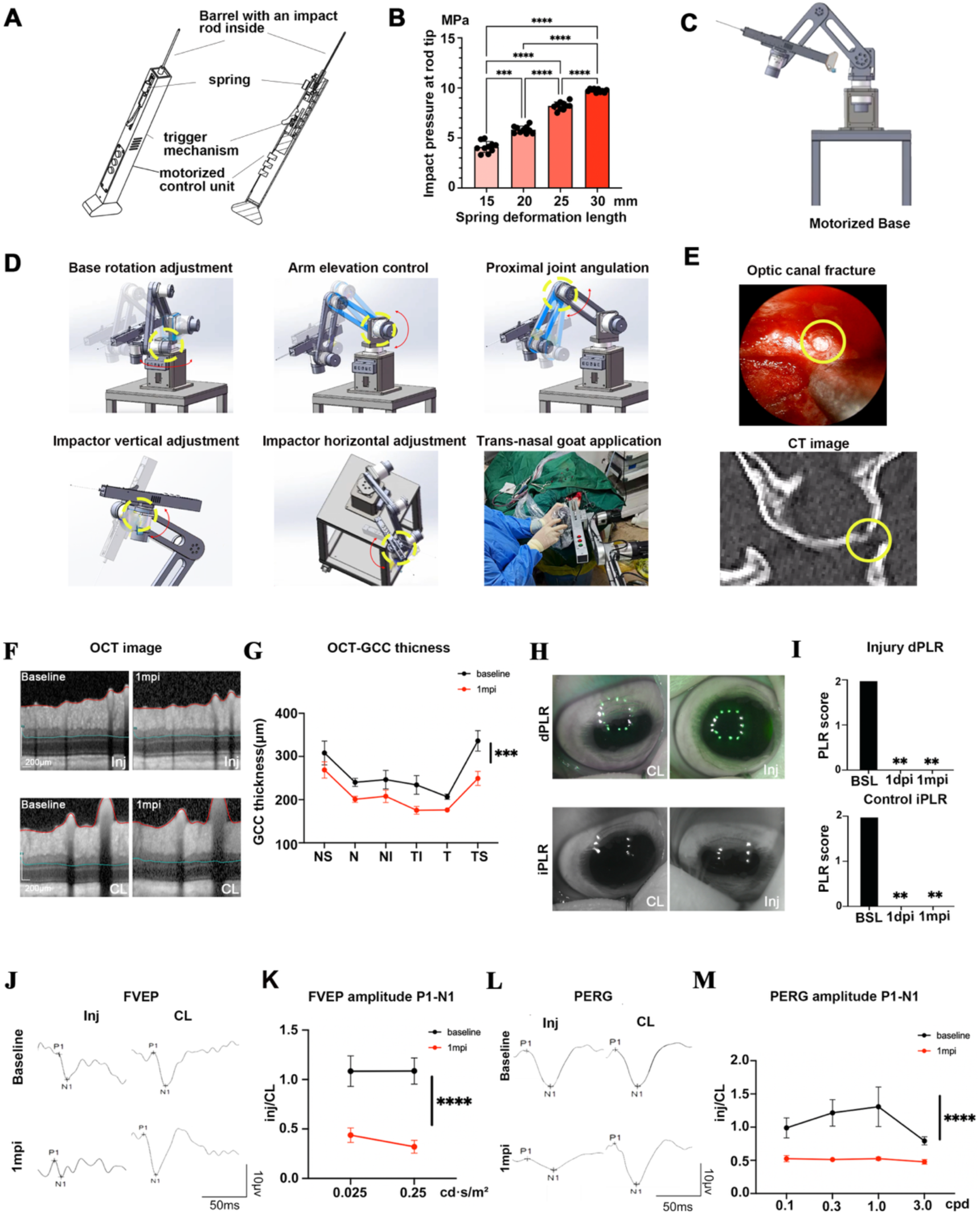
The elastic-energy driven goat model of traumatic optic neuropathy. **(A)** Schematic diagram of the elastic-energy driven impact device used for optic canal fracture induction. **(B)** Histogram summarizing the relationship between spring extension distance and impact force at the rod tip. Each bar represents one extension distance (n = 10 impacts per distance). **(C)** Schematic of the motorized base with five independently driven joints, providing multi-axis positioning and recoil compensation during impact. **(D)** Multi-axis stabilizing base for precise positioning of the impactor. Schematic views illustrate base rotation, arm elevation, proximal joint angulation, and vertical and horizontal adjustment of the impact head, which together allow flexible alignment of the impact rod with the optic canal. The right-bottom panel shows intraoperative transnasal application of the impact system in a goat. **(E)** Verification of impacted optic canal fracture. Representative intraoperative endoscopic view (top) showing a circular bony fragment (yellow circle) that has been driven into and embedded within the optic canal, forming an impacted fracture. Postoperative CT image (bottom) confirms the impacted optic canal fracture, with the bony fragment lodged in the canal lumen (yellow circle) and exerting focal compression on the optic nerve. **(F)** Representative OCT images of the ganglion cell complex (GCC) in the injured (Inj) and contralateral (CL) eyes at baseline and 1 month post-injury (1 mpi) **(G)** Quantification of peripapillary GCC thickness in six sectors (NS, N, NI, TI, T, TS) at baseline and 1 mpi (n = 6 goats; mean ± SEM). In the injured eyes, GCC thickness decreased from 261.9 ± 20.15 μm at baseline to 213.0 ± 15.68 μm at 1 mpi (∼18.7% reduction; P = 0.00022). **(H)** Representative infrared PLR images showing loss of the direct PLR (dPLR) in the injured eye and preserved indirect PLR (iPLR) in the contralateral eye at 1 dpi, consistent with a relative afferent pupillary defect. **(I)** Quantification of dPLR scores in the injured eye and iPLR scores in the contralateral eye at baseline (BSL), 1 day (1 dpi), and 1 month (1 mpi) post-injury. **(J)** Representative FVEP waveforms recorded from injured and contralateral eyes at baseline and 1 mpi (flash intensities 0.025 and 0.25 cd·s/m²). **(K)** Relative FVEP P1–N1 amplitude ratios (Inj/CL) at the two flash intensities (n = 6; mean ± SEM), showing a reduction from 1.086 ± 0.0007625 at baseline to 0.3792 ± 0.05850 at 1 mpi (∼65.1% reduction; P < 0.0001). **(L)** Representative PERG waveforms from injured and contralateral eyes at baseline and 1 mpi (spatial frequency 0.3 cpd). **(M)** Relative PERG P1–N1 amplitude ratios (Inj/CL) across spatial frequencies (0.1–3.0 cpd; n = 6; mean ± SEM), decreasing from 1.076 ± 0.1156 at baseline to 0.5104 ± 0.01148 at 1 mpi (∼52.6% reduction; P < 0.0001 at 0.3 cpd). Impact-force data in (B) were analyzed using ordinary one-way ANOVA with Tukey’s multiple comparisons test.OCT-based GCC thickness in (G) was analyzed using the Scheirer– Ray–Hare non-parametric two-factor test with retinal sector and time (baseline vs 1 mpi) as factors. PLR data in (I) were analyzed using one-way ANOVA. FVEP amplitude ratios in (K) were analyzed using two-way ANOVA with stimulus frequency and time as factors. PERG amplitude ratios in (M) were analyzed using the Scheirer–Ray–Hare non-parametric two-factor test with spatial frequency and time as factors. Data are mean ± SEM; *p < 0.05, **p < 0.01, ***p < 0.001, ***p < 0.001, ****p < 0.0001.

To further enhance precision and reproducibility in large-animal procedures, a motorized stabilizing base incorporating five independently actuated joints was constructed (Fig. 6C). This configuration effectively counteracted recoil during impact and enabled multi-axis positional adjustment (Fig. 6D), converting the system from a handheld device into a mechanically stabilized platform and substantially improving targeting accuracy.

Following fracture induction with the modified device, transnasal endoscopic examination revealed a circular bony fragment displaced into the optic canal lumen, while corresponding sagittal CT imaging confirmed fracture of the anterior wall of the optic canal at the impact site (Fig. 6E).

Structural assessment by OCT demonstrated that GCC thickness in the injured eyes decreased from 261.9 ± 20.15 μm at baseline to 213.0 ± 15.68 μm at 1 mpi, representing an ∼18.7% reduction (P = 0.00022; Fig. 6F,G), whereas no significant thinning was detected in the contralateral eyes (Fig. S3A). These structural changes were comparable to those observed following gas-driven impact, indicating preserved injury severity. Consistent with structural damage, functional assessments revealed unilateral visual deficits. Both the indirect pupillary light reflex (iPLR) in the injured eye and the direct PLR (dPLR) in the contralateral eye remained intact (Fig. S3B), whereas the dPLR in the injured eye was abolished and that in the contralateral eye was preserved (Fig. 6H,I). Electrophysiological measurements again revealed unilateral functional impairment. In the injured eyes, the FVEP P1–N1 amplitude ratio decreased from 1.086 ± 0.0007625 at baseline to 0.3792 ± 0.05850 at 1 mpi (≈65.1% reduction; P < 0.0001; Fig. 6J,K), and the PERG P1–N1 amplitude ratio declined from 1.076 ± 0.1156 to 0.5104 ± 0.01148 (≈52.6% reduction; P < 0.0001; Fig. 6L,M). In contrast, no significant reductions in FVEP or PERG amplitudes were observed in the contralateral eyes compared with baseline values (Fig. S3C,D), confirming a unilateral deficit without widespread bilateral involvement.

## Discussion

### Why optic canal–centered biomechanics matter for modeling TON

TON is widely recognized as a consequence of blunt cranio-orbital trauma, and clinical and radiological studies have long implicated the optic canal as a frequent site of injury, particularly in cases accompanied by optic canal fracture. However, while the anatomical vulnerability of the optic canal is well appreciated, the precise biomechanical pathways by which external forces are transmitted through the cranio-orbital skeleton and concentrated onto the intracanalicular optic nerve have remained insufficiently characterized. This lack of quantitative biomechanical understanding has, in turn, limited the development of experimental TON models that faithfully reflect the clinical injury context.

In particular, existing TON models have largely been defined by empirical injury paradigms rather than by clinically informed mechanical criteria. As a result, although many models reproduce downstream pathological outcomes, such as RGCs loss or visual dysfunction, the biomechanical fidelity of injury induction, especially with respect to optic canal involvement, has been difficult to assess in a standardized and quantitative manner. Consequently, a gap remains between clinically observed optic canal–centered injury patterns and the experimental frameworks used to model TON.

In this study, we addressed this gap by reconstructing a high-resolution human head finite element model to quantitatively characterize force transmission during clinically relevant blunt impacts. Our analyses precisely mapped stress propagation and demonstrated preferential convergence of mechanical stress within the intracanalicular segment across multiple impact directions. These simulations provided a mechanistic basis for defining optic canal–centered injury as a biomechanical target.

Importantly, these finite element–derived insights directly informed the design of our experimental model. We determined that direct optic canal impact represents a biofidelic injury paradigm for TON. Implemented under transnasal endoscopic guidance in goats, this approach enables controlled induction of optic canal fracture and intracanalicular nerve compression while preserving the surrounding ocular structures. In doing so, the model establishes not only anatomical and pathological similarity to human TON, but also a quantitative link between biomechanics and experimental injury induction, providing a more rigorous framework for evaluating disease severity and therapeutic intervention.

From a structural mechanics perspective, the intracanalicular segment is uniquely susceptible to load convergence because it is constrained within a rigid bony canal and surrounded by tightly apposed meningeal and periosteal interfaces (Pircher et al., 2017), leaving limited capacity for displacement or stress redistribution during cranio-orbital impact. Under blunt trauma, forces transmitted along the orbital walls can therefore be funneled toward the optic canal and translated into localized compression and shear on the intracanalicular nerve. This anatomical confinement provides a plausible mechanistic explanation for why the peak loading in our simulations consistently emerged at the intracanalicular segment across multiple impact directions. Importantly, this biomechanical information helps connect experimental TON model to a clinically relevant injury pattern and offers quantitative parameters that can be used to standardize injury severity across studies.

### Comparison with existing TON models

Most existing experimental TON models have been developed in rodents and rely on direct optic nerve crush using forceps or aneurysm clips (Tang et al., 2011). Although these approaches reliably induce RGCs loss and visual pathway dysfunction, they bypass the optic canal entirely and therefore do not recapitulate the canal fracture–mediated injury mechanism characteristic of human TON. In addition, optic nerve crush injuries are typically applied to the retrobulbar segment of the nerve, in close proximity to the globe. This injury location results in rapid retrograde degeneration and early RGCs apoptosis, which differs from the clinical scenario in TON, where optic canal injury occurs at a greater distance from the eye. As a consequence, the temporal dynamics of RGCs degeneration and downstream visual dysfunction in crush models may not accurately reflect the progression observed in TON patients. Together, these anatomical and temporal differences limit the extent to which optic nerve crush models can reproduce clinically relevant patterns of optic canal fracture and disease evolution.

In addition to crush paradigms, more invasive rodent models have also been reported that require cranial opening to access the visual pathway. For example, in a previously described model (Bei et al., 2016), injury is induced by optic tract transection performed following craniotomy, enabling direct and highly controlled disruption of post-chiasmatic visual projections under visual guidance. Although such approaches permit precise manipulation of injury severity, they require extensive cranial opening and direct access to the intracranial optic nerve, resulting in substantial surgical trauma and systemic physiological stress.

More recently, ultrasound-based mouse models have been proposed (Tao et al., 2017), in which suprabrow ultrasonic energy is used to induce TON. These paradigms partially mimic external blunt impact but necessarily deposit acoustic energy in the overlying orbital and periocular soft tissues, often causing collateral damage to extraocular muscles, eyelids, or surrounding bone. Moreover, the short, species-specific anatomy of the murine optic canal and orbit differs substantially from that of humans, which may alter the way mechanical energy is transmitted to the nerve. Thus, both direct crush and ultrasound-mediated rodent models capture important aspects of optic nerve injury but do not fully reproduce the combination of optic canal fracture, localized intracanalicular compression, and preserved contralateral eye that characterizes many human TON cases.

In contrast, the present goat model was designed with an explicit focus on human-relevant biomechanics rather than empirical injury paradigms. Grounded in human-head finite element analysis and implemented through a minimally invasive transnasal approach, this strategy preserves the optic canal–centered injury mechanism. Beyond injury mechanism, goats offer important anatomical and translational advantages. Their cranio-orbital architecture and optic canal geometry more closely resemble human characteristics than rodent models, enabling more realistic force transmission and improved biomechanical fidelity. Importantly, the relatively larger local surgical space surrounding the optic canal in goats allows endoscopic manipulation, device deployment, and interventional refinement, providing opportunities for therapeutic and surgical technology development that are not feasible in murine models. Overall, these features position the goat optic canal fracture model as a biofidelic platform that bridges the gap between small-animal studies and human disease, with clear advantages for mechanistic investigation and preclinical evaluation.

### Advantages of the optimized elastic-energy driven impactor

Accurate and reproducible induction of optic canal injury is a prerequisite for large-animal modeling of TON. In the present study, we refined an initial gas-driven impact system into a second-generation elastic-energy driven device to overcome limitations inherent to pneumatic actuation. Although the gas-driven impactor was instrumental in validating the feasibility of transnasal endoscopic access to the optic canal and impact-induced visual dysfunction, its performance was constrained by several technical and practical factors that limited its suitability for standardized and quantitative studies.

First, operation under high-pressure pneumatic conditions imposed stringent requirements on system sealing and component durability; repeated use was associated with component wear, occasional air leakage, and increased maintenance demands, collectively reducing long-term operational robustness. Second, gas leakage from the storage cylinder led to fluctuations in output force, thereby compromising the consistency of optic canal fracture induction. Third, substantial mechanical recoil during force release frequently resulted in positional deviation of the impact site, limiting spatial accuracy. Fourth, reliance on handheld operation constrained surgical precision and increased susceptibility to operator-dependent variability.

The optimized elastic energy–driven system directly addresses these limitations by replacing gas propulsion with motor-controlled spring extension, enabling precise calibration of impact energy and highly consistent force delivery across animals. Integration with a multi-joint motorized stabilizing base further enhances positional control, minimizes recoil-induced displacement, and effectively eliminates operator-dependent variability. Collectively, these refinements establish a robust and scalable impact platform that supports reproducible optic canal fracture induction and facilitates its application across different surgical postures and large-animal species.

### Limitations and outlook

Despite the strengths of the present model, several limitations should be acknowledged. Large-animal experiments require substantial financial investment and logistical support, which may constrain sample sizes and reduce statistical power. Although the elastic energy–driven impact system improves experimental reproducibility, subtle inter-individual differences in cranio-orbital anatomy may still influence local stress distribution and injury severity. Accordingly, the use of age- and body weight–matched animals is recommended to minimize biological variability. In addition, interspecies differences in skull density, optic canal geometry, and connective tissue composition warrant careful validation before extending this model to nonhuman primates. While OCT and electrophysiology provide clinically aligned, non-invasive structural and functional endpoints to support model validation, deeper tissue-level characterization was not the primary focus of the present platform study. Instead, such analyses will be a focus of follow-up mechanistic and interventional studies building on this model, where optic nerve axonal pathology and retinal ganglion cell loss can be quantified, and extended to investigate white matter injury mechanisms.

Nevertheless, the goat optic canal fracture model provides a clinically meaningful platform for translational studies of TON. From a translational perspective, the modular design of the impact system supports its application as a versatile platform for translational drug evaluation and facilitates adaptation to more advanced experimental settings, including integration with robotic microsurgical technologies and neural interface testing. Extension of this approach to nonhuman primates represents a critical step toward translational and eventual clinical application.

## Conclusion

By combining high-resolution human head finite element analysis with iterative engineering refinement, we identified the intracanalicular segment of the optic nerve as the principal biomechanical vulnerability during blunt periorbital trauma and leveraged this insight to establish a biofidelic, transnasal endoscopic optic canal fracture model in goats. This large-animal model reproducibly recapitulates the defining structural and functional features of clinical TON, including optic canal fracture, GCC thinning, relative afferent pupillary defect, and visual electrophysiological deficits, while minimizing collateral injury and preserving the fellow eye. Building on this framework, the goat model and elastic-energy–driven impact system provide a practical platform for future studies on neuroprotection, axonal regeneration, surgical decompression strategies, and the development of translational interventions targeting optic canal–centered traumatic optic neuropathy.

## Data availability

Data and code will be made available on reasonable request.

## Competing interests

The authors declare no competing interests.

## Author contributions

Conceptualization: Y.Z., W.W., Z.Y.; Data curation: Z.Y., Y.Z.; Formal analysis: H.D., T.Y.; Funding acquisition: W.W., Y.Z.; Investigation: Z.Y., Y.Z., H.D., T.Y., Y.C., S.T., B.X., T.X., H.W., S.L., M.L., S.Z., H. C., S.H., J.Z., Y.W., R.Z.; Methodology: Z.Y., T.Y., Y.C., S.T.; Project administration: Y.Z., Z.Y.; Resources: W.W., Y.Z.; Supervision: Y.Z., W.W., J.Y.; Visualization: Z.Y., T.Y.; Writing – original draft: Z.Y., T.Y.; Writing – review & editing: Y.Z., W.W., J.Y..

## Acknowledgements

We gratefully acknowledge the support of the large-animal housing and care platform at Wenzhou Medical University for providing husbandry and veterinary assistance throughout this study. We thank Zhongshi Technology Co., Ltd. for their assistance in designing and manufacturing the customized impact devices used in this study.

## Funding

This work was supported by: the National Key R&D Program of China (Grant No.2022YFA1105500), the Key Science and Technology Program of Wenzhou (Grant No.ZY2022021) and the National Natural Science Foundation of China (Grant No.82471080).

## Figures

**Figure S1.**
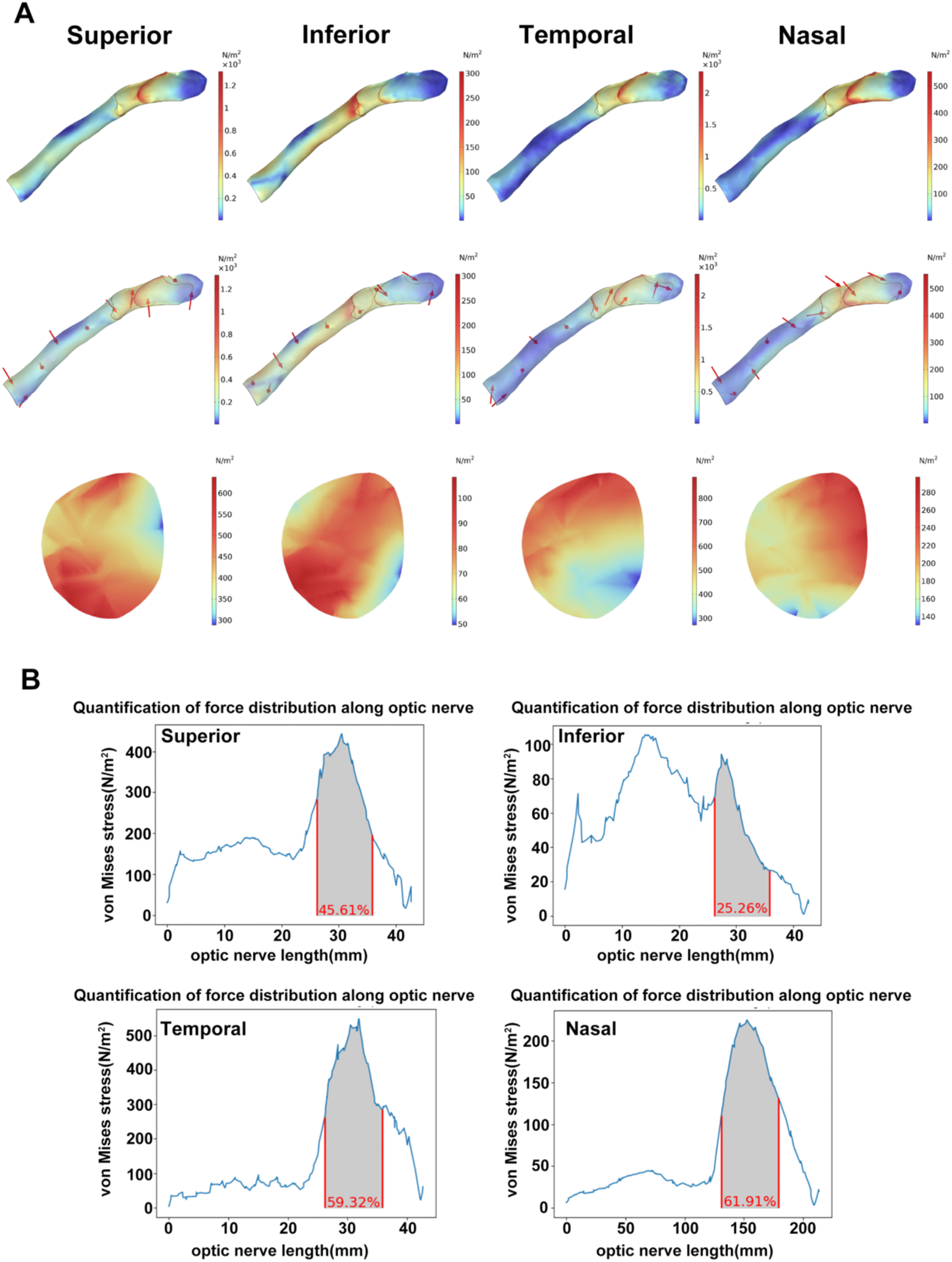
Comparative analysis of force distribution in the optic nerve following periorbital impacts at different locations. **(A)** Illustrations of force distribution along the optic nerve (upper panel), directions of force along the optic nerve (middle panel), and force distribution through the cross-section of the mid intracanalicular optic nerve (lower panel) at 50.6 ms, when the periorbital impact reached its maximum strength following periorbital impacts at various locations (Superior, Inferior, Temporal, Nasal). **(B)** Schematic depicting the locations of periorbital impacts and quantification of force distribution along the optic nerve at 50.6 ms following periorbital impacts at various locations.

**Figure S2.**
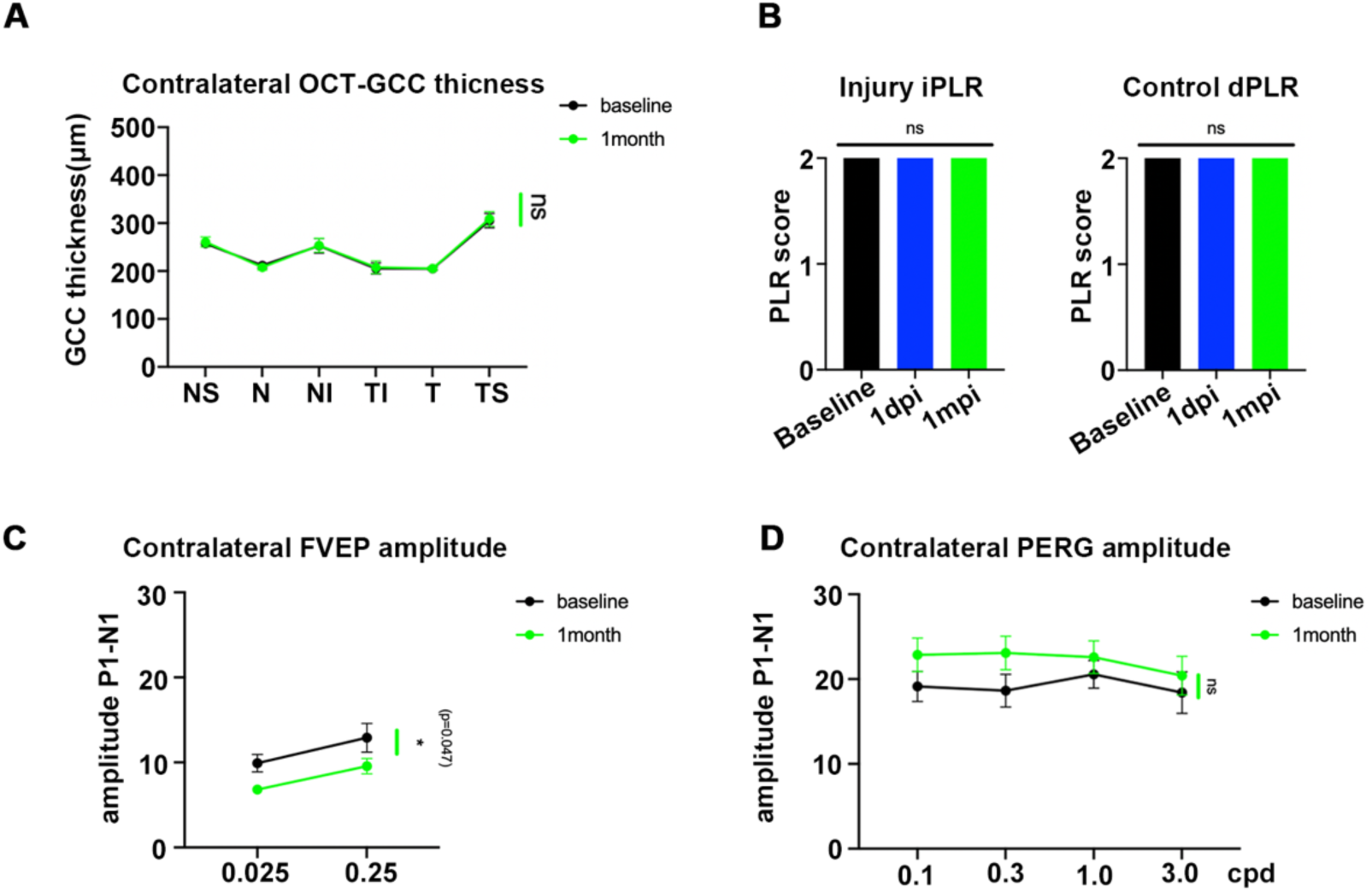
Visual function and GCC thickness are preserved in the contralateral eyes after TON induction with a gas-driven device. **(A–D)** Gas-driven impact device cohort (n = 8 goats). **(A)** OCT-derived GCC thickness in different macular sectors of the contralateral eyes at baseline and 1 mpi (NS, nasal-superior; NI, nasal-inferior; TI, temporal-inferior; T, temporal; TS, temporal-superior).**(B)** PLR scores in the TON cohort showing preserved indirect PLR (iPLR) in the injured eyes and direct PLR (dPLR) in the contralateral eyes at baseline, 1 day post-injury (1 dpi) and 1 mpi (0–2 grading scale; ns). **(C)** FVEP P1–N1 amplitudes of the contralateral eyes at two flash intensities (0.025 and 0.25 cd·s/m²) at baseline and 1 mpi. **(D)** PERG P1–N1 amplitudes recorded from the contralateral eyes at different spatial frequencies (0.1, 0.3, 1.0, and 3.0 cpd) at baseline and 1 mpi. Data are presented as mean ± SEM. OCT, PLR, FVEP and PERG data in (A–D) were analyzed using two-way ANOVA with appropriate post hoc multiple comparisons. ns, not significant.

**Figure S3.**
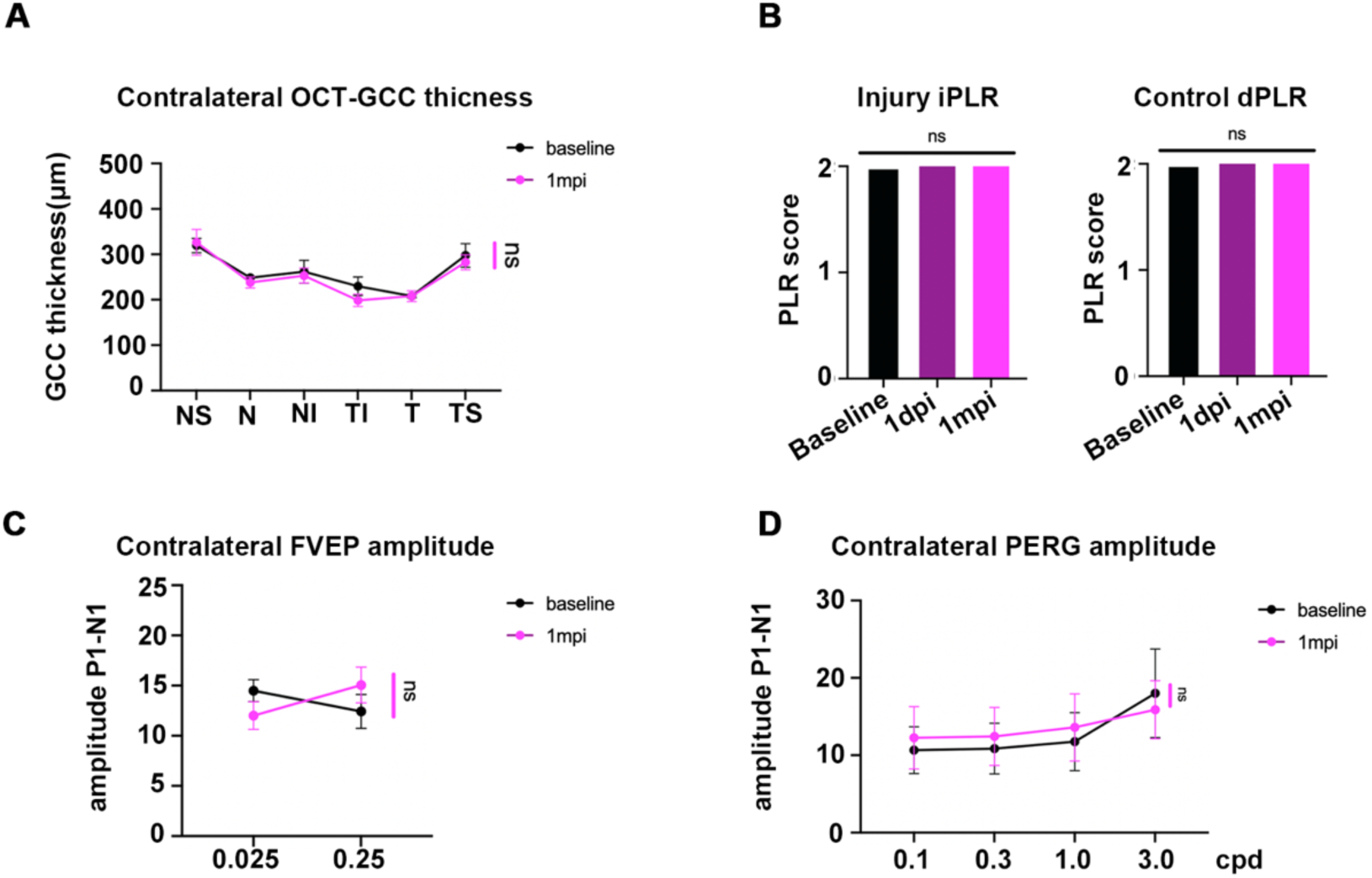
Visual function and GCC thickness are preserved in the contralateral eyes after TON induction with an elastic-energy driven device. **(A–D)** Elastic-energy driven impact device cohort (n = 6 goats). **(A)** OCT-derived GCC thickness in the contralateral eyes at baseline and 1 mpi, **(B)** PLR scores in the TON cohort showing preserved indirect PLR (iPLR) in the injured eyes and direct PLR (dPLR) in the contralateral eyes at baseline, 1 day post-injury (1 dpi) and 1 mpi (0–2 grading scale; ns). **(C)** FVEP P1–N1 amplitudes, and **(D)** PERG P1–N1 amplitudes. Data are presented as mean ± SEM. OCT, PLR, FVEP and PERG data in (A–D) were analyzed using two-way ANOVA with appropriate post hoc multiple comparisons. ns, not significant.

## Notes

### Competing Interest Statement

The authors have declared no competing interest.

